# Multidimensional semantic representations emerge from multi-frequency representational similarity learning

**DOI:** 10.64898/2026.09.23.753859

**Authors:** Saskia L. Frisby, Christopher R. Cox, Ajay D. Halai, Akihiro Shimotake, Takayuki Kikuchi, Katsuya Kobayashi, Takeharu Kuneida, Yoshiki Arakawa, Akio Ikeda, Riki Matsumoto, Timothy T. Rogers, Matthew A. Lambon Ralph

## Abstract

Current theories of semantic representation are agnostic about how fine-grained neurophysiological activity corresponds to representation of multidimensional semantic information. By applying an innovative multivariate technique (representational similarity learning; RSL), we adjudicated three hypotheses: (1) that multidimensional semantic structure is represented within a single frequency range (e.g., gamma/high gamma); (2) that each semantic dimension is independently represented within a different frequency range; and (3) that multidimensional semantic information is “transfrequency” (at least some information emerges only when multiple frequencies are considered together). RSL was applied to time-frequency power and phase data extracted from electrocorticography (ECoG) grid electrodes on the surface of human ventral anterior temporal lobe (vATL). We found significant decoding of graded, multidimensional semantic information from a wide range of frequencies (4 – 200 Hz), but not from individual frequency bands, providing clear evidence that multidimensional semantic information is coded in a transfrequency fashion.

## 1. Introduction

Semantic cognition is our ability to understand objects, words and events and to engage in meaningful action. The ventral anterior temporal lobes (vATLs) are known to support this ability (Rogers et al., 2004; Patterson et al., 2007; Lambon Ralph et al., 2017) by encoding semantic information that is *graded* (robins, wrens and ostriches are all birds, but the robin and the wren are more similar to each other than either is to the ostrich) and *multidimensional* (robins, wrens and ostriches all have feathers and beaks but differ in colour, size and flying abilities; Lambon Ralph and Patterson, 2008; Lambon Ralph et al., 2010; Cox et al., 2024). However, it is currently unknown how these representational feats are achieved via neurophysiological activity within in the vATL (Murphy, 2024). To address this question, we applied a new analytic approach, representational similarity learning (RSL), to a large sample of electrocorticography (ECoG) data (n = 18) to investigate which frequency ranges contain graded, multidimensional semantic information. By doing so, we adjudicated three competing hypotheses about how vATL activity corresponds to semantic representation.

Considerable previous research has established that semantic representation is supported by multiple “spokes”, each encoding modality-specific information, plus a bilateral vATL “hub” that synthesises information across modalities and temporal instances and thereby enables generalisation across contexts (Rogers et al., 2004; Patterson et al., 2007; Lambon Ralph et al., 2010, 2017; Jackson et al., 2021). This theory is supported by convergent evidence from multiple sources: neuropsychological studies of patients with vATL atrophy (Hodges et al., 1992; Bozeat et al., 2000; Hodges and Patterson, 2007), positron emission tomography (PET; Devlin et al., 2000), distortion-corrected functional magnetic resonance imaging (Binney et al., 2010; Visser et al., 2010; Halai et al., 2014); magnetoencephalography (MEG; Mollo et al., 2017), neuro-computational simulations (Rogers et al., 2004, 2021; Chen et al., 2017), transcranial magnetic stimulation (TMS; Pobric et al., 2007, 2010a, 2010b), and invasive cortical stimulation mapping (Lüders et al., 1991; Shimotake et al., 2015; Matoba et al., 2024). However, these sources of evidence primarily serve to establish *that* the vATL is necessary for full semantic representation, rather than *how* the vATL activity encodes multidimensional representations. The precise relationship between semantic information and the power and phase of neural activity is unclear (Murphy et al., 2026). Multiple kinds of correspondence are possible:

1. One or a few frequency ranges carry information about graded, multidimensional semantic structure. Several previous studies have associated semantic tasks with activity in a specific frequency band, although they differ in which band they implicate – theta (Hermes et al., 2014), alpha (Clarke et al., 2018), beta (Abel et al., 2015; Sato et al., 2021), gamma, or high gamma (Crone et al., 2001; Crone and Hao, 2002; Tanji, 2005; Edwards et al., 2010; Cervenka et al., 2011; Chan et al., 2011; Wang et al., 2011; Kojima et al., 2013; Nakai et al., 2017, 2019; Forseth et al., 2018; Arya, 2019; Bartoli et al., 2019; Snyder et al., 2023). This first hypothesis would predict that graded, multidimensional semantic structure is decodable from one or a limited number of frequency ranges. Note that frequency ranges exhibit interdependence (e.g. theta-gamma coupling; Sederberg et al., 2003; Lisman, 2005; Canolty et al., 2006; Canolty and Knight, 2010; Hermes et al., 2014; Aru et al., 2015; Heusser et al., 2016; Benítez-Burraco and Murphy, 2019), and so, if this hypothesis is true, some coarse semantic information, e.g. animacy, may still be decodable from frequency ranges that are not used as the primary means of representation (“weak” representation; (Watrous et al., 2015; Frisby et al., 2026b).
2. Each dimension is encoded independently by a separate frequency range. This second hypothesis would predict that one graded dimension is decodable from each frequency range, that the dimension decoded differs between frequency range, and that multidimensional structure is evident when decoders are trained on all ranges simultaneously.
3. Multidimensional semantic information is “transfrequency”, by which we mean that at least some information is encoded jointly by multiple frequency ranges and is not evident when those ranges are considered independently. We (Frisby et al., 2026b) previously found that regularised logistic regression classifiers trained to decode animacy based on power within individual frequency bands (theta, alpha, beta, gamma, and high gamma) produced significant decoding accuracy, but classifiers trained on power from the whole frequency spectrum (4 – 200 Hz) produced significantly higher decoding accuracy. However, animacy is a coarse distinction and so whether this is true of graded, multidimensional semantic information is an open question. This third hypothesis would predict that multiple dimensions can be decoded *only* when all frequency ranges are considered together.

Elucidation of the neural code within the vATL requires the novel combination of two crucial opportunities: (1) direct recordings of neurophysiological activity in the vATL during semantic tasks and (2) methods capable of revealing the detailed correspondence between that vATL activity and graded, multidimensional semantic information. The first of these requirements is met by human ECoG data - ECoG is a direct, spatially resolved (∼ 1 cm) and temporally precise (∼ 1 ms) technique that, unlike noninvasive techniques such as electroencephalography (EEG), is sensitive to a wide range of frequencies including very high-frequency activity. The second is met by representational similarity learning (RSL; (Cox et al., 2024)), a cutting-edge multivariate decoding method that uses neural data to predict coordinates within a multidimensional target space. Unlike encoding methods that have previously been applied to ECoG time-frequency data (Wang et al., 2011; Rupp et al., 2017), RSL makes very few assumptions about the nature of the underlying neural code. The few assumptions that are made can be fine-tuned by the experimenter in order to align with the experimenter’s explicit hypotheses about how information is likely to be represented – for example, the assumption that information is likely to be encoded by relatively small populations of neurons with correlated activity (Cox and Rogers, 2021; Frisby et al., 2023, 2026a; Cox et al., 2024; see Methods). Additionally, unlike representational similarity analysis (RSA; (Kriegeskorte et al., 2008b, 2008a)), RSL calculates correlations for each dimension of the target information separately, making it a powerful tool for differentiating between the three hypotheses described above. A previous study demonstrated that RSL can successfully reveal semantic information in ECoG data (Cox et al., 2024), but that study analysed voltage, which reflects a conflation of high- and low-frequency activity, leaving the key hypotheses of this study unexplored.

We adjudicated these three hypotheses by applying RSL to power and phase extracted from ECoG data collected as 18 participants named pictures. To test our first and second hypotheses, we applied RSL to power and phase data within a single frequency range – theta (4 – 7 Hz), alpha (8 - 12 Hz), beta (13 – 30 Hz), gamma (30 – 60 Hz) and high gamma (60 – 200 Hz). To test our third hypothesis, we applied RSL to power and phase data from the whole frequency range (4 – 200 Hz).

## 2. Methods

These data were previously described by Frisby et al. (2026b). The data and the preprocessing pipeline are described in detail there and are summarised here.

### 2.1. Patients

Nineteen patients participated in the study (labelled 01-22 to facilitate comparison with previous work; Shimotake et al., 2015; Chen et al., 2016; Rogers et al., 2021; Cox et al., 2024; Frisby et al., 2026b). One patient was excluded because too many trials were contaminated with artefacts, leaving 18 patients. All patients were native speakers of Japanese. Table 1 contains information about patients’ age, sex, handedness, and clinical presentation.

**Table 1:** Patient characteristics. Abbreviations: WAIS-R – Wechsler Adult Intelligence Scale (1991), WAIS-III - Wechsler Adult Intelligence Scale (1997), VIQ – Verbal IQ, PIQ – Performance IQ, TIQ – full- scale IQ, WMS-R – Wechsler Memory Scale (1987), WAB – Western Aphasia Battery, FAS – focal aware seizure, FIAS – focal impaired awareness seizure, FBTCS – focal to bilateral tonic-clonic seizure, aMTG – anterior middle temporal gyrus, PHG – parahippocampal gyrus, pMTG – posterior middle temporal gyrus, mITG – medial inferior temporal gyrus, SMG - supramarginal gyrus, ITG – inferior temporal gyrus, IPL – intraparietal lobule, HS – hippocampal sclerosis, HA – hippocampal atrophy, FCD – focal cortical dysplasia. * - The WAIS-R was used to test patients 01-06 and the WAIS-III was used to test other patients. ** - missing score.

|  | Patient 01 | Patient 02 | Patient 03 | Patient 04 |
| --- | --- | --- | --- | --- |
| Age, sex, handedness | 22, M, R | 29, M, R&L | 17, F, R | 38, F, R |
| WAIS-R/WAIS-III* (VIQ, PIQ, TIQ) | 70, 78, 69 | 72, 78, 72 | 67, 76, 69 | 84, 97, 89 |
| WMS-R (verbal, visual, general, attention, delayed recall) | 99, 64, 87, 91, 82 | 99, 92, 97, 87, 83 | 51, <50, <50, 81, 56 | 75, 111, 83, 62, 53 |
| WAB | 95.6 | 96 | 97.2 | 98.5 |
| WADA | Left | Bilateral | Left | Left |
| Age of seizure onset | 16 | 10 | 12 | 29 |
| Seizure type | FAS→FIAS, FBTCS | FAS→FIAS | FAS→FIAS | FAS→FIAS |
| Ictal ECoG onset | aMTG | PHG | PHG | PHG |
| MRI | L basal frontal cortical dysplasia, L anterior temporal arachnoid cyst | L posterior temporal cortical atrophy | L temporal tip arachnoid cyst | L HS/HA |
| Pathology | FCD type I | FCD type IIIa | Palmini FCD type IB | FCD type IIa |
|  | Patient 05 | Patient 06 | Patient 07 | Patient 08 |
| Age, sex, handedness | 55, M, R | 34, M, L | 41, F, R | 27, F, R |
| WAIS-R/WAIS-III* (VIQ, PIQ, TIQ) | 105, 99, 103 | 55, **, 44 | 72, 83, 75 | 106, **, 105 |
| WMS-R (verbal, visual, general, attention, delayed recall) | 71, 117, 84, 109, 72 | 52, <50, <50, 55, <50 | 83, 111, 89, 94, 82 | 112, 114, 114, 81, 100 |
| WAB | 98 | 88 | 97.3 | 99.6 |
| WADA | Left | Left | Right | Left |
| Age of seizure onset | 55 | 12 | 19 | 16 |
| Seizure type | FIAS (once) | FAS→FIAS | FAS→FIAS | FAS→FIAS |
| Ictal ECoG onset | None | R parietal lobe/pMTG | PHG | ventral anterior temporal |
| MRI | L medial temporal lobe low-grade glioma | R parietal cerebral atrophy & contusion, R hippocampal sclerosis/atrophy | L HS/HA, L parieto-occipital perinatal infarction | R medial temporal cyst |

Table 1: Patient characteristics.
| Pathology | Diffuse astrocytoma | Post-traumatic change(parietal)/scar(temporal)/HS | FCD type I | FCD type I |
| --- | --- | --- | --- | --- |
|  | Patient 09 | Patient 10 | Patient 11 | Patient 12 |
| Age, sex, handedness | 51, M, R | 38, F, R | 29, F, R | 40, M, L |
| WAIS-R/WAIS-III* (VIQ, PIQ, TIQ) | 73, 97, 83 | 109, 115, 112 | 62, 80, 67 | 93, 105, 98 |
| WMS-R (verbal, visual, general, attention, delayed recall) | 80, 101, 85, 91, 91 | 71, 79, 70, 90, 58 | 64, 94, 68, 79, 79 | 74, 94, 77, 110, 96 |
| WAB | 89.6 | 96.9 | 95.8 | 99 |
| WADA | Left | Left | Left | Right |
| Age of seizure onset | 43 | 28 | 12 | 6 |
| Seizure type | FIAS | FAS→FIAS | FAS→FIAS | FAS→FIAS |
| Ictal ECoG onset | mITG | SMG | PHG | PHG |
| MRI | L temporal cavernoma | L parietal operculum tumour | L HS | R HS/HA |
| Pathology | Arteriovenous malformation | Oligoastrocytoma | non-neoplastic brain tissue | FCD type IIIa |
|  | Patient 13 | Patient 14 | Patient 15 | Patient 20 |
| Age, sex, handedness | 22, M, R | 42, M, R | 35, M, R | 23, F, R |
| WAIS-R/WAIS-III* (VIQ, PIQ, TIQ) | 86, 79, 81 | 96, 84, 90 | 82, 86, 82 | 67, 82, 71 |
| WMS-R (verbal, visual, general, attention, delayed recall) | 55, 79, 53, 90, 54 | 73, 85, 73, 103, 76 | 75, 89, 75, 92, 81 | 78, 119, 86, 82, 64 |
| WAB | 97.4 | 99.9 | 99.2 | 94.2 |
| WADA | Left | Left | Left | Bilateral |
| Age of seizure onset | 14 | 27 | 20 | 15 |
| Seizure type | FAS→FIAS | FAS→FIAS | FIAS | FIAS |
| Ictal ECoG onset | PHG | PHG | PHG | IPL/SMG |
| MRI | L HS/HA | L HS | L HS/HA | No apparent lesion |
| Pathology | FCD type IIIa | FCD type IIIa | FCD type IIIa | FCD type I |
|  | Patient 21 | Patient 22 |  |  |
| Age, sex, handedness | 40, M, R | 28, F, R |  |  |
| WAIS-R/WAIS-III* (VIQ, PIQ, TIQ) | 86, 97, 90 | 80, 69, 72 |  |  |
| WMS-R (verbal, visual, general, attention, delayed recall) | 111, 113, 113, 87, 109 | 77, 89, 77, 121, 73 |  |  |
| WAB | 92.2 | 100 |  |  |
| WADA | Left | Left |  |  |
| Age of seizure onset | 30 | 12 |  |  |
| Seizure type | FIAS | FIAS |  |  |
| Ictal ECoG onset | aMTG | PHG |  |  |
| MRI | No apparent lesion | L HS/HA |  |  |

Patients were implanted with grids or strips of platinum subdural electrodes for presurgical monitoring (recording diameter 2.3 mm, inter-electrode distance 1cm, ADTECH, WI). A clinical MPRAGE T_1_-weighted anatomical scan was acquired before and after electrode implantation to be used for electrode localisation. 15 patients had electrodes implanted in the left hemisphere, of which 10 – 42 electrodes (mean 25 electrodes) were within grids or strips that covered the vATL (where grids or strips spanned both ventral anterior temporal cortex and posterior or lateral temporal cortex, all electrodes in the grid or strip were included in the analysis). The remaining 3 patients had electrodes implanted in the right hemisphere, of which 6 – 28 electrodes (mean 20 electrodes) were within grids or strips that covered the vATL. Figure 1 shows the locations of the electrodes.

**Figure 1:**
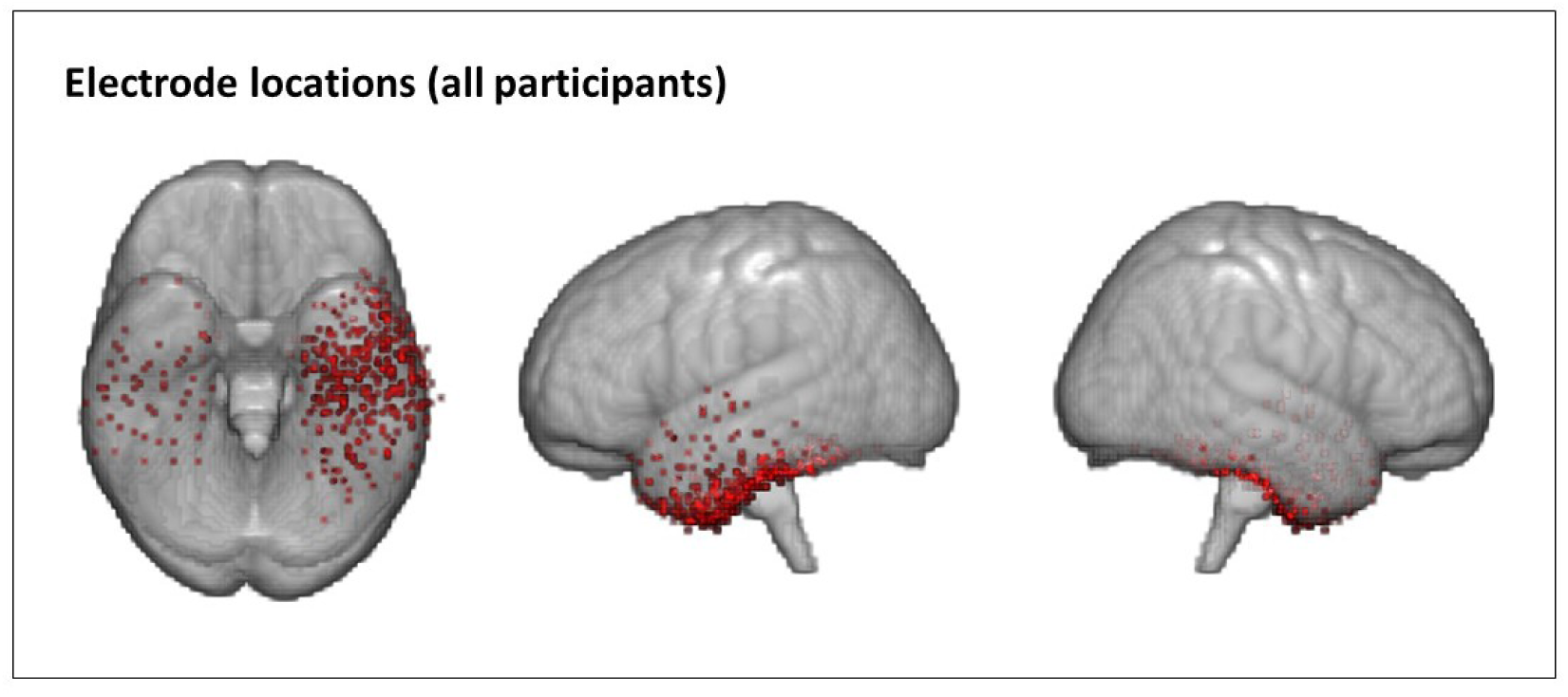
Electrode locations for all patients. Electrodes are overlaid on an MNI template (MNI152NLin2009cAsym).

All patients gave written informed consent and the study was approved by the ethics committee of the Kyoto University Graduate School of Medicine (#C533).

### 2.2. Stimuli, task and acquisition

Stimuli were 100 line drawings – 50 animals and 50 nonliving items. Items in the two categories did not differ in low-level visual structure, visual complexity, concreteness, familiarity, word frequency, or age of acquisition (Barry et al., 1997; Morrison et al., 1997; Shimotake et al., 2015; Chen et al., 2016; Rogers et al., 2021; Cox et al., 2024; Frisby et al., 2026b). The stimuli were displayed on a PC screen using MATLAB r2010a.

Each patient completed four runs of the task in a single session. Within each run, each stimulus appeared once, in a random order. Each stimulus was presented for five seconds and there was no break between stimuli. Patients were instructed to name each picture as quickly and accurately as possible. Time of naming onset was recorded and eye fixation was monitored via video.

Data for nine patients were recorded at 2000 Hz (with a low-pass filter of 600 Hz) and data for ten patients were recorded at 1000 Hz (with a low-pass filter of 300 Hz).

### 2.3. Data analysis

#### 2.3.1. Preprocessing – structural MRI

The position of each electrode was identified on each 2D slice of the post-surgical scan. Next, electrode positions were coregistered to the pre-surgical scan and then to MNI space using *fnirt* (https://fsl.fmrib.ox.ac.uk/fsl/fslwiki/; Jenkinson et al., 2012; Smith et al., 2004). The position of each electrode was then manually adjusted to the surface.

#### 2.3.2. Preprocessing – ECoG

For full details of the ECoG preprocessing pipeline, including analysis of the impact of preprocessing on subsequent decoding accuracy, we refer the reader to Frisby et al. (2026b).

In summary, the following steps were implemented in MATLAB r2023b using functions from EEGLAB (2023.1; https://eeglab.org/; Delorme & Makeig, 2004): (1) removing line noise at 60 Hz and the harmonics 120 and 180 Hz using the CleanLine EEGLAB plugin (v2.0; https://github.com/sccn/cleanline/ ; Delorme, 2023; Mitra & Bokil, 2007); (2) removing slow drifts with a high-pass filter of 0.5 Hz; (3) applying a low-pass filter of 300 Hz for consistency across participants (see Stimuli, task and acquisition), (4) rejection of channels below the seizure onset zone or with poor contact; (5) epoching between -1000 and 3000 ms relative to stimulus onset; (6) baseline-correction to the mean response across trials between -200 and -1 ms; (7) downsampling to 1000 Hz, via boxcar averaging, for consistency across participants; (8) common average referencing; and (9) rejection of trials containing artefacts. The artefact rejection stage comprised (9A) automatically rejecting trials containing values more extreme than 10 standard deviations from the mean for that channel; (9B) inspecting all trials and rejecting trials containing interictal activity or “electrode pop”; (9C) applying ICA to extract *n* components where *n* is 75% of the number of good electrodes, then using SASICA (https://github.com/dnacombo/SASICA/; Chaumon et al., 2015) to extract and remove trials containing muscle activity; and (9D) filtering with a saccade-related potential template and counting the number of microsaccades in each component (since every component contained very few microsaccades - < 0.000012 microsaccades per second - no components were rejected at this stage).

Time-frequency power was extracted using complex Morlet wavelet convolution (Bertrand et al., 1994) for every trial between 0 and 1650 ms in 10 ms timesteps and between 4 and 200 Hz in 60 logarithmically-spaced frequency steps. A five-cycle wavelet was used at 4 Hz, increasing to a 15-cycle wavelet at 200 Hz (Clarke, 2020). Power values were then averaged across repeated presentations of the same stimulus and any missing trials were interpolated with the median power across all trials. Decibel normalisation was performed using the mean power across trials (for each electrode and frequency) between -300 and -100 ms. Since phase values cannot be averaged (Cohen, 2014), trials were averaged across repeated presentations of the same stimulus before phase values were extracted using the same wavelet parameters used for extracting power. Voltage values were obtained by averaging preprocessed data over repeated presentations of the same stimulus.

Preprocessing code is available at https://github.com/slfrisby/ECoG_LASSO/.

#### 2.3.3. Multivariate decoding

##### 2.3.3.1. Approach

This section provides an overview of our RSL approach. For mathematical detail, and a thorough comparison of RSL to RSA, we refer the reader to Oswal et al. (2016) and Cox et al. (2024).

RSL models use *input features* (in this case, neural data) to predict the coordinates of items within a multidimensional *target space*. Target spaces are constructed such that, within the target space, items that are hypothesised to be represented more similarly are located closer together. The hypotheses tested in this study were that power or phase within various frequency ranges represents semantic information, so a semantic target space was constructed. This was achieved (1) by taking vectors of feature-rating norms for each concept (Dilkina and Lambon Ralph, 2012), (2) creating a representational similarity matrix (RSM) by calculating the similarity between those vectors for each possible pair of stimuli, and (3) applying singular value decomposition to the RSM to extract 3 components (accounting for 81.1%, 4.4%, and 4.0% of variance in the whole RSM), which were used as the three dimensions of the target space (Cox et al., 2024). As shown in Figure 2, items that are more semantically similar are closer in the target space – therefore, if an RSL model can use time-frequency power and/or phase extracted from the vATL to successfully predict one or more of a stimulus’s coordinates in the target space, this would indicate that power or phase within the vATL encodes semantic information.

**Figure 2:**
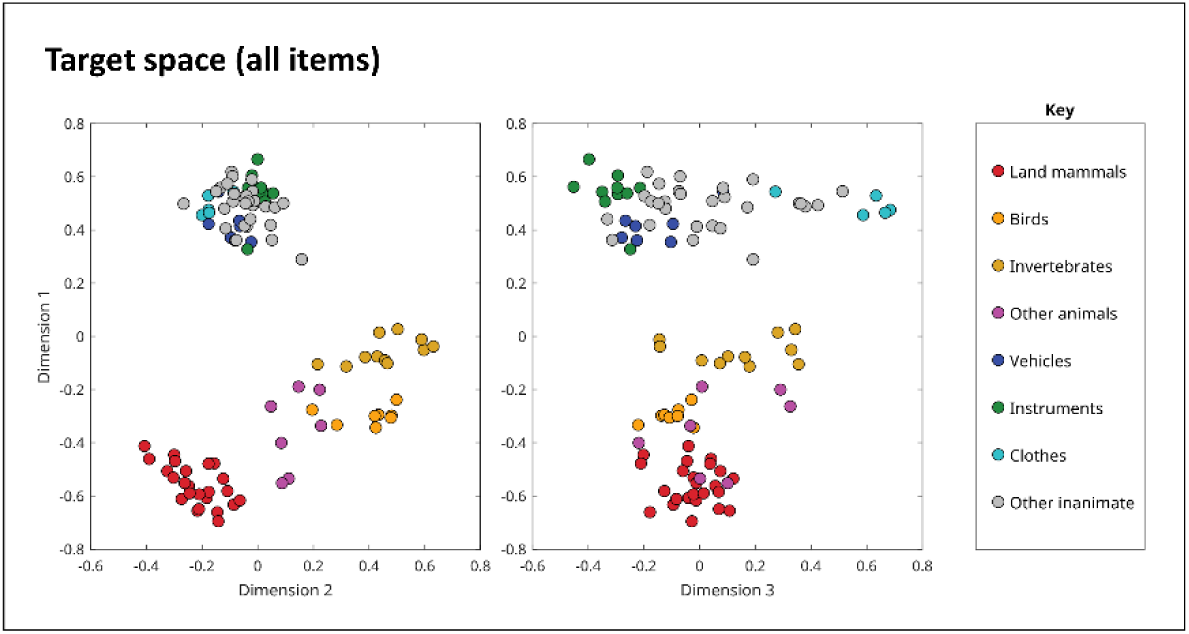
Coordinates of all stimuli on each target semantic dimension. Colours indicate category membership – land mammals (red), birds (orange), invertebrates (mustard), other animals (magenta), vehicles (navy), instruments (green), clothes (turquoise), and other inanimate objects (grey).

Vectors of power or phase input features were created for each item at each timepoint (0 ms, 10 ms, 20 ms…) by concatenating power or phase values for all electrodes in a single participant for all frequencies in a range of interest. Vectors of voltage input features were created by concatenating voltage values for all electrodes in a single participant in a 50 ms window centred on the timepoint of interest. Note that power, phase, and voltage features all reflect activity *around*, as well as at, the timepoint of interest – power and phase features do because Morlet wavelet convolution takes activity at neighbouring timepoints into account, and voltage features do because the window is 50 ms wide.

An RSL model is the matrix formulation of a regularised linear regression model – it predicts coordinates on multiple dimensions by fitting multiple coefficients (one coefficient per dimension) to multiple input features. The models used group-ordered-weighted LASSO (grOWL) regularisation (Oswal et al., 2016), which preferentially creates models in which (1) few of the total features receive nonzero coefficients, (2) features that do receive nonzero coefficients are correlated in their activity, and (3) features that receive *any* nonzero coefficients receive *all* (i.e. three) nonzero coefficients. These constraints can be thought of as three assumptions that models trained with grOWL regularisation make: (1) that the neural code is relatively sparse, (2) that neurons, neural populations, and/or frequencies encoding the target information are correlated in their activity, and (3) that neurons, populations, or frequencies that encode the target information encode all dimensions, not just one. The first two of these assumptions, taken together, mean that grOWL assumes that the target information is encoded by small groups of correlated neurons, populations and/or frequencies. This is our working hypothesis and is arguably more plausible than the alternative claim that only one electrode or frequency encodes our target information in a “grandmother cell” fashion (Quiroga et al., 2008), or the alternative that *all* electrodes and frequencies studied encode the target information, to the exclusion of other information needed for cognitive processes (Frisby et al., 2023). The third preference means that, if it is the case that a single feature encodes more than one dimension, grOWL is likely to discover that feature. Additionally, the presence of single features encoding multiple dimensions would indicate parallels between the vATL and deep layers of computational models. Within these deep layers, the activity patterns of individual units do not correspond to interpretable features such as *has eyes* or *has feathers* – yet activity patterns across multiple units are gradedly more similar for more similar concepts (Rogers and McClelland, 2004; Rogers et al., 2004; Jackson et al., 2021; Giallanza et al., 2025). Note that these assumptions are best thought of as loose preferences, not rigid constraints – for example, if neural features encode only one dimension, then models trained with grOWL can still learn to predict that one dimension well. Models were trained using the WISC MVPA toolbox in MATLAB r2018b (https://github.com/crcox/WISC_MVPA/) using default parameters.

Model performance was assessed using ten-fold nested cross-validation. The stimuli were divided into ten folds of ten stimuli each. One of these folds was designated as the outer-loop holdout set and another was designated as the inner-loop hold-out set. The remaining eight folds were used as training data to search for the best value of the hyperparameter *λ* (which controls the trade-off between minimising prediction error and prioritising the preferences of the regularisation penalty). Models with different hyperparameter values were evaluated on the inner-loop holdout set. The folds were then reassigned so that a different fold was used as the inner-loop holdout set and the previous inner-loop holdout set was used for training. Once all combinations of inner-loop holdout set and training data had been explored, a final model was trained on all folds except the outer-loop holdout set using the best-performing hyperparameter value. This model was then tested on the outer-loop holdout set and the correlation between predicted coordinates and target coordinates was calculated. The whole procedure was repeated with each of the ten folds being used as the outer-loop holdout set. The average of the ten correlations between predicted coordinates and target coordinates was our measure of model performance.

Additionally, the predicted coordinates were visualised and compared to the target coordinates in Figure 2. When models are trained, the targets are standardised with respect to the values in the training set, which means that predicted coordinates for each test set are scaled differently (for further detail, see Cox et al., 2024). This does not affect the calculation of performance, which is done for each fold individually, but it would skew a visualisation of the predicted coordinates. To adjust for this, for each predicted coordinate, we calculated the mean of the training set coordinates used to make that prediction and then added that mean to the predicted coordinate. To visualise change in predicted coordinates over time, coordinates were plotted in MATLAB r2023b, and ffmpeg 2203 (https://ffmpeg.org) was used to concatenate plots at successive timepoints with a frame rate of ten (ten times slower than real time). For ease of visualisation, we created similar plots in which coordinates for each item were averaged within category (land mammals, birds, invertebrates, other animals, vehicles, instruments clothes, and other inanimate items) and animated these averages.

Code for the decoding analysis is available at https://github.com/slfrisby/ECoG_RSL/.

##### 2.3.3.2. Experimental questions

To reiterate, we wished to adjudicate three hypotheses – (1) that gamma and/or high gamma power encodes multiple graded semantic dimensions, but other frequency ranges encode only binary animacy; (2) that different dimensions can be decoded from different frequency ranges; or (3) that representation is transfrequency, i.e., at least some dimensions are encoded jointly by multiple frequency ranges but are not evident when those ranges are considered independently.

###### 2.3.3.2.1. What information is evident in individual frequency ranges?

To test the first two hypotheses, we divided the 60 frequencies into theta (4 – 7 Hz, 11 frequencies), alpha (8 - 12 Hz, 7 frequencies), beta (13 – 30 Hz, 13 frequencies), gamma (30 – 60 Hz, 10 frequencies) and high gamma (60 – 200 Hz, 19 frequencies) ranges. We created vectors of power or phase input features for each item at each timepoint, using only frequencies within a single range. We trained models using these features (separately for each participant) and generated a timecourse of mean correlation between predicted and target coordinates. We averaged these timecourses over participants and compared each group-average timecourse to zero using one-tailed, one-sample t-tests. Probabilities were adjusted to control the false-discovery rate at α = 0.05 (Benjamini and Hochberg, 1995).

Significant correlations on more than one dimension would indicate that the frequency range encodes multidimensional information. However, those correlations are not conclusive evidence for graded structure. As shown in Figure 2, dimension 1 separates animate from inanimate items, but also distinguishes between items within each domain. A model that predicted the same low coordinate for every animate item and the same high coordinate for every inanimate item (i.e. a model that predicted a category, but not a graded dimension) would produce significant correlations with the target. Within-domain, however, the model’s predictions would not correlate significantly with the target. Therefore, to verify that representations are graded, we calculated correlations just for animate items and just for inanimate items.

###### 2.3.3.2.2. What information is evident in frequency ranges considered together?

To test the third hypothesis, we created vectors of power or phase features using all frequencies from 4 to 200 Hz and used these to train models. We also wished to investigate how these models performed relative to models trained on voltage, which reflects a conflation of a wide range of frequencies. Therefore, we also created voltage feature vectors, which included values within a 50 ms window centred on the timepoint of interest (see Approach). Again, we calculated correlations for all items, for animate items, and for inanimate items, and we compared each group-average timecourse to zero using one-tailed, one-sample t-tests, adjusting probabilities to control the false-discovery rate at α = 0.05. We compared power and voltage, and phase and voltage, using paired t-tests, controlling the false-discovery rate.

## 3. Results

### 3.1. What information is evident in individual frequency ranges?

To assess the first two hypotheses – (1) that gamma and/or high gamma power encodes multiple graded semantic dimensions, but other frequency ranges encode only binary animacy; (2) that different dimensions can be decoded from different frequency ranges – we trained RSL models to predict each item’s coordinates within a three-dimensional target space derived from semantic feature-rating norms (Dilkina and Lambon Ralph, 2012; Cox et al., 2024). As shown in Figure 2, dimension 1 of this space separates animate from inanimate items and also differentiates subcategories of animal, dimension 2 further differentiates animals, and dimension 3 primarily differentiates inanimate items.

These models were trained on power or phase features from a single frequency range –theta (4 – 7 Hz), alpha (8 - 12 Hz), beta (13 – 30 Hz), gamma (30 – 60 Hz) and high gamma (60 – 200 Hz). Figure 3 shows the results. Models trained on power within any range could successfully predict dimension 1, but all models failed to predict dimensions 2 and 3. Models trained on phase with the theta range could predict dimension 1, but models trained on any other range could not. This result is incompatible with our first hypothesis (that multidimensional semantic structure is represented within a single frequency range). Since the same dimension 1 was decodable from each frequency range, this result is also compatible with our second hypothesis (that each semantic dimension is independently represented within a different frequency range).

**Figure 3:**
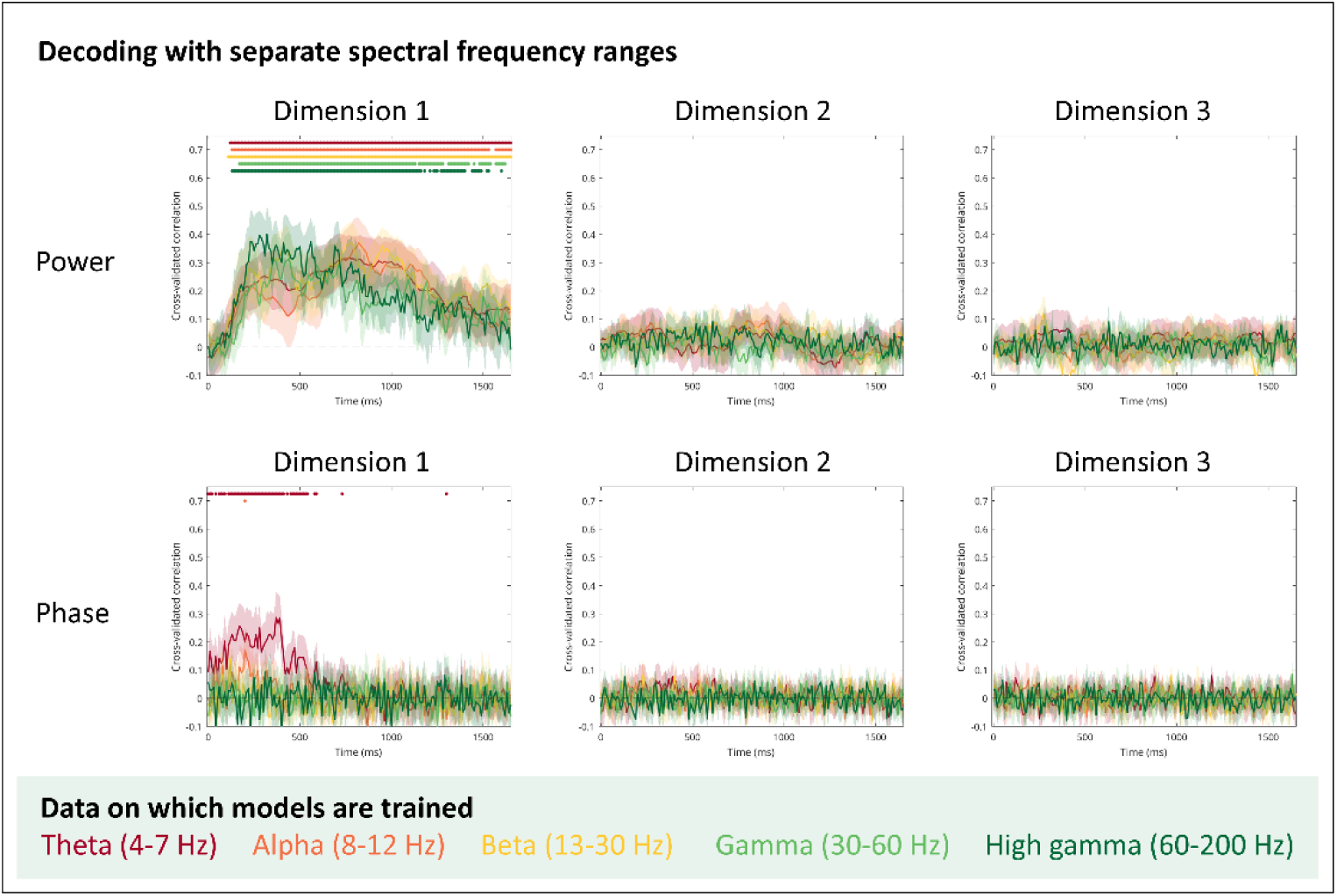
Decoding power and phase within individual frequency ranges. (A) Mean and 95% confidence interval of the hold-out correlations for RSL models trained to predict coordinates on three semantic dimensions based on power frequency features from a given range -– theta (4–7 Hz, red), alpha (8–12 Hz, orange), beta (13–30 Hz, yellow), gamma (30–60 Hz, light green), and high gamma (60–200 Hz, dark green). Coloured dots indicate a significant difference between classifier accuracy and chance (0.5, one-sample t-tests with probabilities adjusted to control the false-discovery rate at α = 0.05). (B) Mean and 95% confidence interval of the hold-out correlations for RSL models trained on phase frequency features from a given range.

To explore this result further, we evaluated cross-validated correlations within animate and inanimate domains separately, which arise only when significant correlations reflect truly graded structure and not when they reflect binary encoding of animacy (see Methods). Figure 4A shows that, for power, within-domain correlations were significant for animate stimuli within all ranges and for inanimate stimuli within the alpha range only. These results indicate that all frequency ranges encode a single dimension, but that the dimension is graded, not only distinguishing between animate and inanimate items but also capturing variation within-domain. By contrast, Figure 4B shows that, for phase, within-domain correlations were not significant for theta phase (or for any other frequency band). Therefore, we found no evidence that theta phase represented any more than a binary animacy distinction.

**Figure 4:**
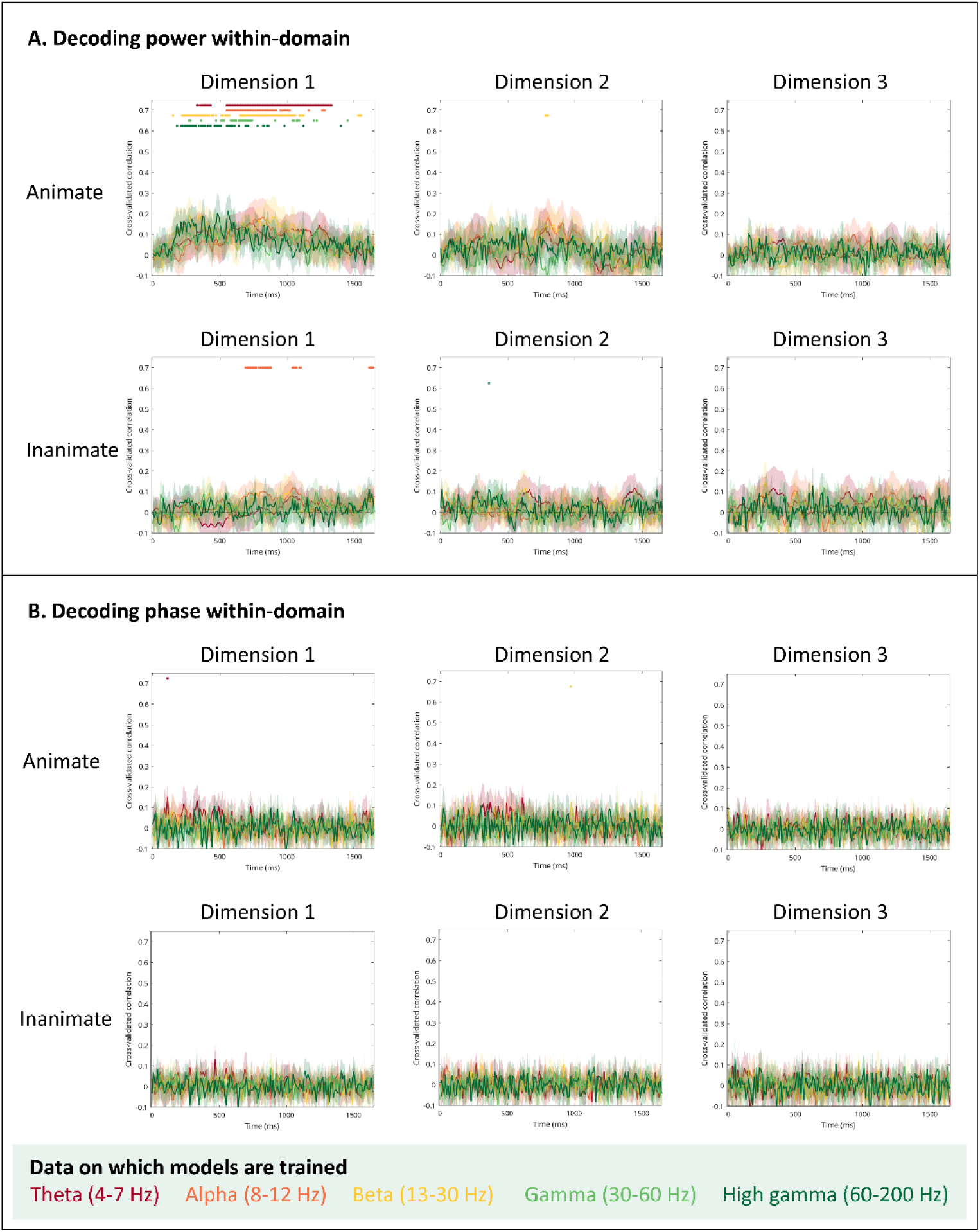
Decoding performance within-domain. A) Mean and 95% confidence interval of the hold-out correlations, calculated for animate or for inanimate stimuli only, for RSL models trained to predict coordinates on three semantic dimensions based on power frequency features from a given range -– theta (4–7 Hz, red), alpha (8–12 Hz, orange), beta (13–30 Hz, yellow), gamma (30–60 Hz, light green), and high gamma (60–200 Hz, dark green). Coloured dots indicate a significant difference between classifier accuracy and chance (0.5, one-sample t-tests with probabilities adjusted to control the false-discovery rate at α = 0.05). (B) Mean and 95% confidence interval of the hold-out correlations, calculated for animate or for inanimate stimuli only, for RSL models trained on phase frequency features from a given range.

### 3.2. What information is evident in frequency ranges considered together?

Next, we tested the third hypothesis - that representation is transfrequency, meaning that at least some dimensions are encoded jointly by multiple frequency ranges but are not evident when those ranges are considered independently. We trained models on power or phase features using all frequencies from 4 to 200 Hz and, as a comparison, on voltage features extracted from a 50 ms window centred on the timepoint of interest. Figure 5 shows the results. Models trained on power from all frequency ranges could successfully predict the first two dimensions, both for all items and within-domain. Similarly, models trained on voltage, which reflects the combination of power within different frequency ranges, could also predict the first two dimensions for all items and within-domain. Differences between models trained on these two kinds of data were not significant. Figure 5 also shows results for phase. These models could predict the first dimension, but were always considerably outperformed by models trained on voltage. These results constitute clear novel evidence in support of our third hypothesis – that representation of graded, multidimensional semantic information by power is transfrequency.

**Figure 5:**
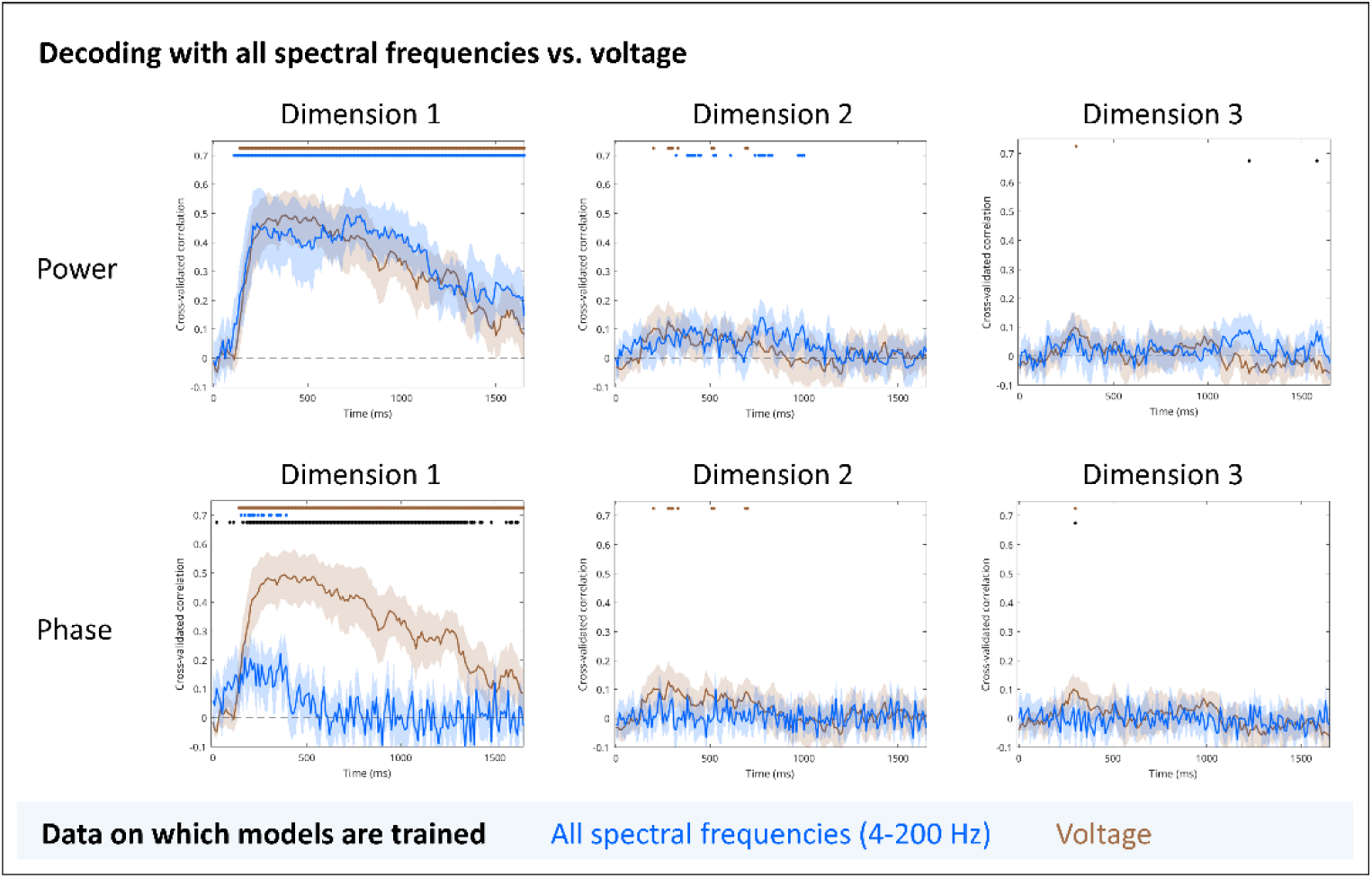
Decoding power and phase from all frequencies. (A) Mean and 95% confidence interval of the hold-out correlations for RSL models trained to predict coordinates on three semantic dimensions based on power frequency features from all frequencies (4 – 200 Hz, blue) or from voltage features (brown). Coloured dots indicate a significant difference between classifier accuracy and chance (0.5, one-sample t-tests with probabilities adjusted to control the false-discovery rate at α = 0.05). Black dots indicate a significant difference between accuracies at a given timepoint (paired t-tests with probabilities adjusted to control the false-discovery rate at α = 0.05). (B) Mean and 95% confidence interval of the hold-out correlations for RSL models trained on phase frequency features from all frequencies (4 – 200 Hz) or from voltage.

Figure 6 shows cross-validated correlations calculated separately for animate and inanimate domains. Figure 6A shows that, for power, within-domain correlations were significant for animate stimuli for both the first and second dimensions, providing evidence for graded representation. By contrast, Figure 6B shows that, for phase, no within-domain correlations were significant. Consistent with the findings for individual frequency bands, we found no evidence that phase encoded anything more than a binary animacy distinction.

**Figure 6:**
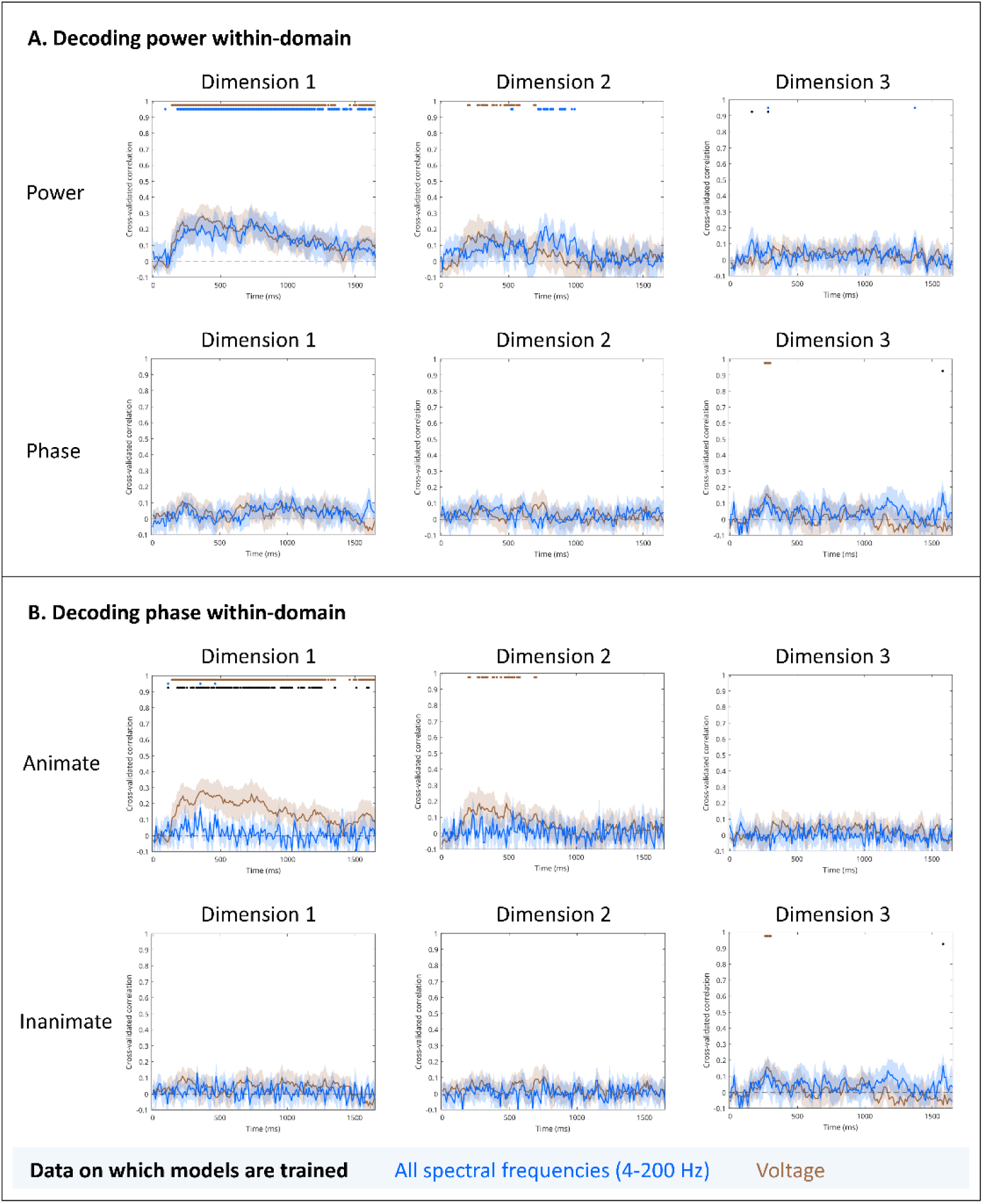
Decoding performance within-domain. (A) Mean and 95% confidence interval of the hold-out correlations, calculated for animate or for inanimate stimuli only, for RSL models trained to predict coordinates on three semantic dimensions based on power frequency features from all frequencies (4 – 200 Hz, blue) or from voltage features (brown). Coloured dots indicate a significant difference between classifier accuracy and chance (0.5, one-sample t-tests with probabilities adjusted to control the false-discovery rate at α = 0.05). Black dots indicate a significant difference between accuracies at a given timepoint (paired t-tests with probabilities adjusted to control the false-discovery rate at α = 0.05). (B) Mean and 95% confidence interval of the hold-out correlations, calculated for animate or for inanimate stimuli only, for RSL models trained on phase frequency features from all frequencies (4 – 200 Hz) or from voltage.

To gain a more detailed understanding of this representation, we visualised the three-dimensional coordinates that the models predicted. We focused on models that could predict coordinates on multiple dimensions, i.e., those trained on power and those trained on voltage (see https://github.com/slfrisby/ECoG_RSL for results for models trained on just one frequency range).

For ease of visualisation, we averaged the model’s predicted coordinates over items within the same category (land mammals, birds, invertebrates, other animals, vehicles, instruments clothes, and other inanimate items). Put another way, each category “centroid” can be considered to reflect predicted coordinates for a “prototypical” category member (Mervis and Rosch, 1981). Figure 7 shows the target coordinates averaged over items within the same category. (We also plotted predicted coordinates without averaging within categories; these results are available at https://github.com/slfrisby/ECoG_RSL).

**Figure 7:**
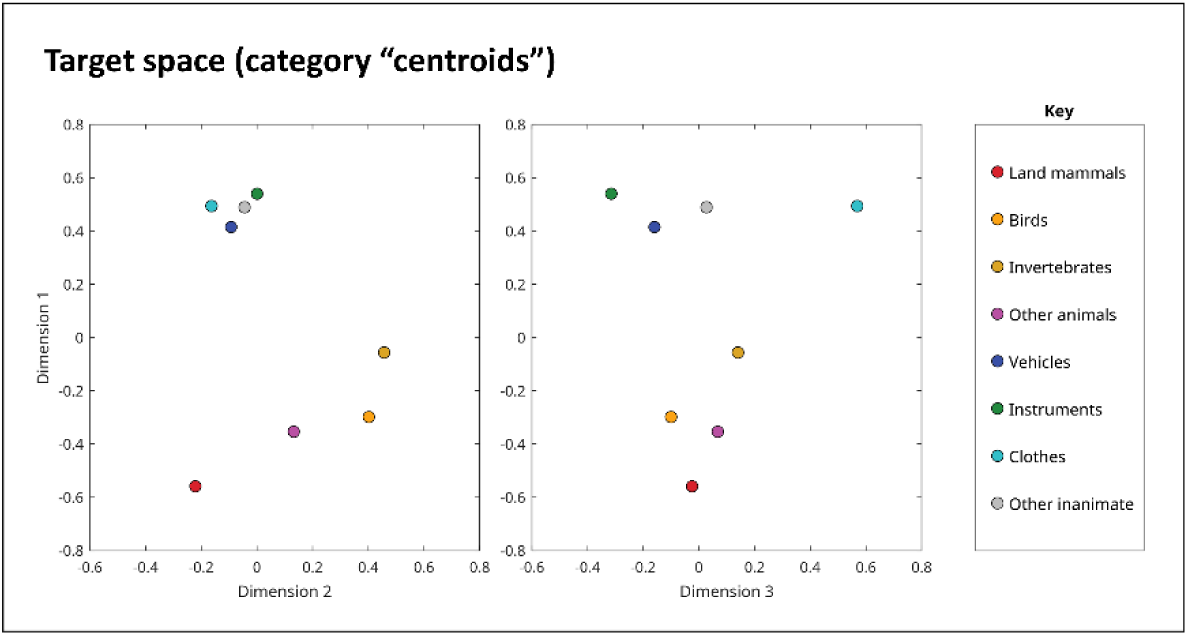
Coordinates of stimuli, averaged over categories, on each target semantic dimension – land mammals (red), birds (orange), invertebrates (mustard), other animals (magenta), vehicles (navy), instruments (green), clothes (turquoise), and other inanimate objects (grey).

Results for models trained on power are shown in Figure 8 and in Video 1. Immediately after stimulus onset, when no information can be decoded, centroids cluster close to the origin. At about 200 ms, centroids swarm away from the origin, which is reflected in a rise in decoding accuracy (Video X). Notably, it is *not* the case that animate categories travel together and inanimate categories travel together before progressive differentiation within-domain. Instead, we observe that, as soon as centroids leave the origin, centroids for animate categories also spread out along dimension 1 – land mammals take the most extreme values, followed by birds, “other” animals (including a bat, a whale, and a frog), and finally invertebrates (see Figure 7). This is reflected in the fact that within-domain decoding follows the same timecourse as decoding of all items (Figures 5 and 6 and Video 1). After their initial flight, centroids for animate categories hover for a while within largely non-overlapping territories. The trajectories of inanimate categories are more overlapping, consistent with the finding that dimension 1 cannot be decoded within the inanimate domain (Figure 6 and Video 1).

**Figure 8:**
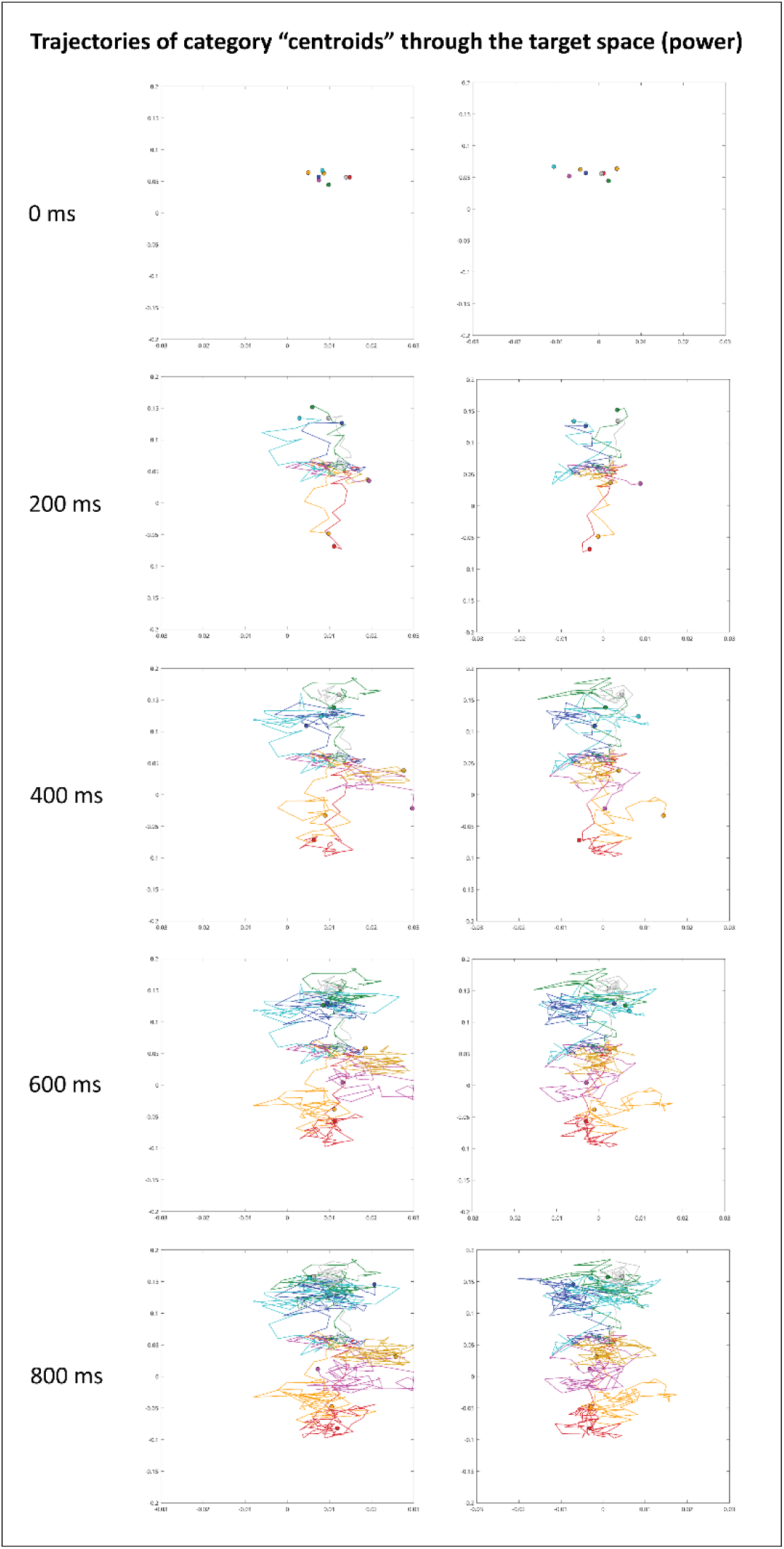
Trajectory of predicted coordinates through target space. RSL models were trained to predict coordinates on three semantic dimensions based on power frequency features from all frequencies (4 – 200 Hz). Coloured dots show coordinates of stimuli, averaged over categories, on each target semantic dimension – land mammals (red), birds (orange), invertebrates (mustard), other animals (magenta), vehicles (navy), instruments (green), clothes (turquoise), and other inanimate objects (grey). Lines show the “trail”, i.e. the previously occupied coordinates, of each dot.

Figure 9 and Video 2 show that models trained on voltage exhibit the same pattern. (Similar animations for individual frequency ranges and for phase can be found at https://github.com/slfrisby/ECoG_RSL).

**Figure 9:**
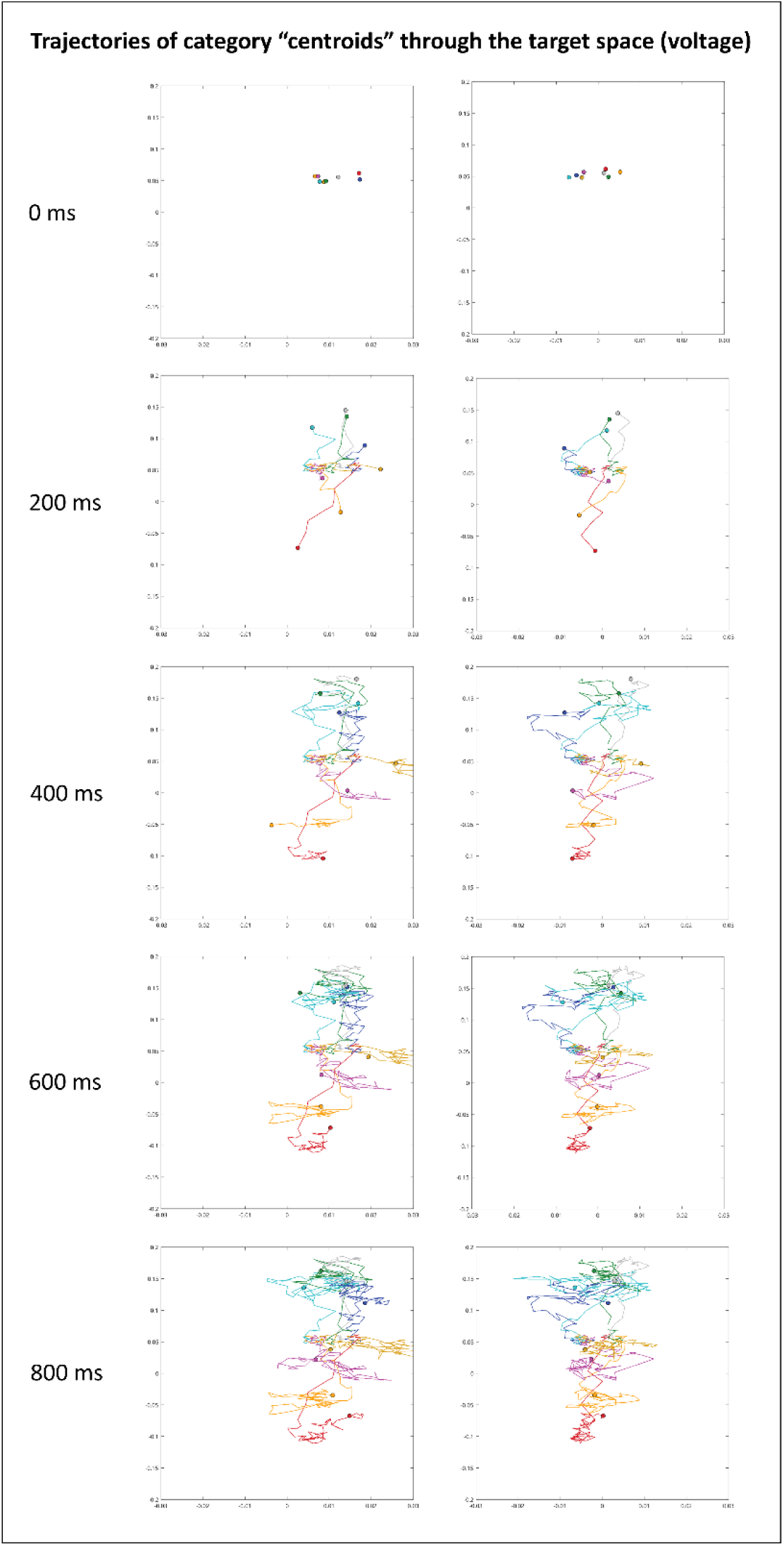
Trajectory of predicted coordinates through target space. RSL models were trained to predict coordinates on three semantic dimensions based on voltage features. Coloured dots show coordinates of stimuli, averaged over categories, on each target semantic dimension – land mammals (red), birds (orange), invertebrates (mustard), other animals (magenta), vehicles (navy), instruments (green), clothes (turquoise), and other inanimate objects (grey). Lines show the “trail”, i.e. the previously occupied coordinates, of each dot.

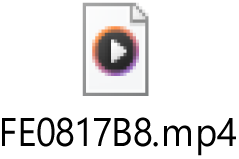

Video 1 (click to play): Trajectory of predicted coordinates through target space. RSL models were trained to predict coordinates on three semantic dimensions based on power frequency features from all frequencies (4 – 200 Hz). The line plot shows the mean hold-out correlations, calculated for all animals for dimension 1, animate items for dimension 2, and inanimate items for dimension 3. Coloured dots above the line plot indicate a significant difference between classifier accuracy and chance (0.5, one-sample t-tests with probabilities adjusted to control the false-discovery rate at α = 0.05). In the plots below, coloured dots show coordinates of stimuli, averaged over categories, on each target semantic dimension – land mammals (red), birds (orange), invertebrates (mustard), other animals (magenta), vehicles (navy), instruments (green), clothes (turquoise), and other inanimate objects (grey). Lines connected to each dot show the “trail”, i.e. the previously occupied coordinates, of each dot.

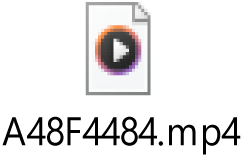

Video 2 (click to play): Trajectory of predicted coordinates through target space. RSL models were trained to predict coordinates on three semantic dimensions based on voltage features. The line plot shows the mean hold-out correlations, calculated for all animals for dimension 1, animate items for dimension 2, and inanimate items for dimension 3. Coloured dots above the line plot indicate a significant difference between classifier accuracy and chance (0.5, one-sample t-tests with probabilities adjusted to control the false-discovery rate at α = 0.05). In the plots below, coloured dots show coordinates of stimuli, averaged over categories, on each target semantic dimension – land mammals (red), birds (orange), invertebrates (mustard), other animals (magenta), vehicles (navy), instruments (green), clothes (turquoise), and other inanimate objects (grey). Lines connected to each dot show the “trail”, i.e. the previously occupied coordinates, of each dot.

## 4. Discussion

Although it is well-established that the vATLs are necessary for the representation of semantic information (Rogers et al., 2004; Patterson et al., 2007; Lambon Ralph et al., 2017), it is currently unknown how graded, multidimensional representations are coded by neurophysiological activity within the vATL. In this study, we found (1) that multidimensional semantic structure cannot be decoded from a single frequency range, and (2) that it is not possible to decode different semantic dimensions from different ranges. Instead, we established that (3) multidimensional semantic information is revealed only when activity from multiple frequency ranges is given as input to the decoder.

Characterisation of *how* (not simply *that*) the vATL represents semantic information depends on the fusion of two crucial opportunities: (1) direct recordings of neurophysiological activity in the human vATL during semantic tasks, and (2) methods capable of revealing the detailed correspondence between that vATL activity and graded, multidimensional semantic information. By decoding fine-grained semantic information from multiple sets of features (time-frequency within or across ranges), we revealed a pattern of results that does not merely support one hypothesis, but rules out others. We believe that leveraging patterns of multivariate results to adjudicate hypotheses in this way – a strategy that we term the *comparative multivariate approach* (Frisby et al., 2026a) – will become a powerful tool for resolving conflicting patterns of results in cognitive neuroscience.

Our findings supported the conclusions that representations within the vATL are transfrequency. Why might this be so? A pervasive perspective is that frequency is inversely related to transmission distance (Canolty and Knight, 2010; Kopell et al., 2010; Lachaux et al., 2012).

Proponents of the hub-and-spoke model (Rogers et al., 2004; Patterson et al., 2007; Lambon Ralph et al., 2017) argue that both vATL hub and unimodal spokes are necessary for semantic representation, so transfrequency properties may reflect both local processing and communication between hub and spokes. Note that the frequency-inverse-to-distance assumption was originally based on the claim that individual neurons cannot fire at speeds within the high-gamma range, and so high-gamma activity must reflect the asynchronous firing of neurons directly below the electrode. This assumption is known to be violated in the case of auditory nerves that can individually produce high-gamma activity (Carlyon et al., 2010; Sevgili, 2023), raising the possibility that cortical neurons could do the same. Additionally, low-frequency activity, thought to reflect long-range connections, has been observed in excised brain tissue (Florez et al., 2015). This demonstrates that the frequency-inverse-to-distance assumption is not symmetrical – even if it is true that cortical neurons cannot individually produce high-gamma activity and so high gamma reflects asynchronous local firing, lower frequencies could reflect local as well as distal activity. Perhaps more progress can be made by discarding assumptions about distance and regarding the “transfrequency” code simply as a voltage code (cf. (Miller, 2010).

We identified two minor exceptions to the overall pattern evident in our results. Firstly, we could decode dimension 1 from theta phase at about 0 – 500 ms post stimulus onset. Given that decoding of theta phase within-domain was not significant (Figure 4), this suggests that theta phase represents a binary animacy distinction rather than any graded semantic structure. Theta phase has been proposed to play various roles, including abstraction of transtemporal concepts from multiple episodic experiences (Sauseng and Klimesch, 2008), encoding of semantic similarity in a manner analogous to the encoding of physical space (Solomon et al., 2019; Spiers, 2020), or retrieving and integrating semantic information in a context-appropriate way (Halgren et al., 2015; Marko et al., 2019; Jackson, 2021). It is not clear how any of these functions could be underpinned by a binary animacy code. A more plausible explanation is theta-gamma coupling – phase-locking of gamma power to theta phase (Sederberg et al., 2003; Canolty et al., 2006; Canolty and Knight, 2010; Lisman and Jensen, 2013; Aru et al., 2015). Correlation of theta phase with gamma and/or high gamma power, from which graded semantic information can be decoded, could explain why theta phase comes to capture at least part of that structure, even if this “weak” representation is not used downstream (Watrous et al., 2015). Future studies should aim to test this idea directly (Proix et al., 2022).

Secondly, we could briefly decode dimension 1 from alpha power within the inanimate domain, which was not the case for the other frequency ranges nor for whole-spectrum decoding. Recall that our aim in evaluating models within-domain was to test for gradedness. In this respect, this additional finding does not lead us to a different conclusion – alpha power represents dimension 1 in a graded way, just as the other frequency ranges do. It is nevertheless interesting that detailed information about inanimate stimuli should be captured by alpha power, especially since dimension 1 does not separate out the inanimate stimuli to any great extent (Figure 1). Proposed roles for alpha in semantic representation include organisation of incoming visual input (Clarke et al., 2018) and “semantic orientation” – the ability to integrate, and orient oneself within, the meaning of objects in the environment (Klimesch, 2012). However, it is not immediately obvious how representation of a single semantic dimension (albeit a graded one) could facilitate either of these processes.

Finally, visualising the trajectories of items through the semantic space during the course of processing reveals some interesting phenomena that future theories of semantic representation should incorporate. Computational models of semantic representation demonstrate a pattern of progressive differentiation during learning – the model first learns superordinate distinctions (such as animacy), then fine-grained detail. Rogers and Patterson (2007) posited that an analogous process takes place on-the-fly during stimulus processing. For example, when seeing a pigeon, the hub’s activation patterns were thought to gradually home in on the “pigeon” region of semantic space, which is located within the broader “animal” region. Therefore, if pressured to respond quickly in the context of a speeded categorisation task, participants would be faster to verify that a stimulus is an “animal” than that it is a “pigeon”, and this was what Rogers and Patterson (2007) found at the behavioural level. However, this distinct sequential pattern is not what we found in our neural data – category “centroids” head towards their territories in a “fountaining” pattern, separating within-domain as soon as they have separated across-domain (Videos 1 and 2). This pattern is unaccounted-for by current theoretical formulations of the hub-and-spoke model (Lambon Ralph et al., 2017) and future iterations of the theory should seek to do so. Note that “fountaining” is also observed in the hub of computational models of semantic cognition (Rogers et al., 2021), which means that we are already ideally situated to explore the possible mechanistic roles of this pattern *in silico*.

## 5. Conclusion

In this study we aimed to adjudicate three hypotheses: (1) that multidimensional semantic structure is represented within a single frequency range (e.g. gamma/high gamma); (2) that each semantic dimension is independently represented within a different frequency range; and (3) that multidimensional semantic information is “transfrequency” (at least some information emerges only when multiple frequencies are considered together). These results provide evidence against the first and second hypotheses and evidence in favour of the third. As well as representing an important step towards a complete characterisation of semantic representation, these results reveal novel representational patterns that future iterations of the hub-and-spoke model should seek to characterise in detail.

## Data and Code Availability

We are unable to share raw data for this study because the patients did not provide informed consent to do so. However, matrices containing power, phase, and voltage features (columns) for each stimulus (rows) are available at https://osf.io/m5v42/. Preprocessing code is available at https://github.com/slfrisby/ECoG_LASSO/ and decoding code is available at https://github.com/slfrisby/ECoG_RSL/.

## Author Contributions

Saskia L. Frisby – Conceptualisation, Methodology, Formal Analysis, Data Curation, Writing – Original Draft, Writing – Review & Editing, and Visualisation. Christopher R. Cox – Conceptualisation, Writing – Review & Editing. Ajay D. Halai – Conceptualisation, Methodology, Formal Analysis, and Writing – Review & Editing. Akihiro Shimotake – Investigation, Data Curation, Writing – Review & Editing, and Supervision. Takayuki Kikuchi – Investigation, Data Curation, and Methodology. Takeharu Kuneida – Investigation, Data Curation, and Methodology. Yoshiki Arakawa – Investigation, Data Curation, and Methodology. Ryosuke Takahashi – Investigation. Akio Ikeda – Investigation, Data Curation, and Methodology. Riki Matsumoto – Investigation, Methodology, and Writing – Review & Editing. Timothy T. Rogers – Conceptualisation, Methodology, Writing – Review & Editing, Supervision, and Funding Acquisition. Matthew A. Lambon Ralph – Conceptualisation, Methodology, Writing – Review & Editing, Supervision, and Funding Acquisition.

## Acknowledgments

This work was supported by an MRC Career Development Award (MR/V031481/1) to A.D.H., by the Japan Agency for Medical Research and Development (25K02548), by the Japan Society for the Promotion of Science (KAKENHI 22H02945, 23KK0146, 25K02548) to R.M., and by an Advanced European Research Council (ERC) award (GAP 670428-30 BRAIN2MIND_NEUROCOMP), MRC programme grant (MR/R023883/1), and intramural funding (MC_UU_00005/18) to M.A.L.R. We would like to thank the patients who so selflessly participated in this study. We would also like to thank the Research Facilitators at the Centre for High-Throughput Computing, University of Wisconsin-Madison, who provided invaluable assistance with the high-throughput computational implementation of RSL.

## Open Access Statement

For the purpose of open access, the authors have applied a Creative Commons Attribution (CC BY) licence to any Author Accepted Manuscript arising from this work.

## References

Abel TJ, Rhone AE, Nourski KV, Kawasaki H, Oya H, Griffiths TD, Howard MA, Tranel D (2015) Direct physiologic evidence of a heteromodal convergence region for proper naming in human left anterior temporal lobe. J Neurosci 35:1513–1520.

Aru J, Aru J, Priesemann V, Wibral M, Lana L, Pipa G, Singer W, Vicente R (2015) Untangling cross-frequency coupling in neuroscience. Curr Opin Neurobiol 31:51–61.

Arya R (2019) Similarity of spatiotemporal dynamics of language-related ECoG high-gamma modulation in Japanese and English speakers. Clin Neurophysiol Off J Int Fed Clin Neurophysiol 130:1403–1404.

Barry C, Morrison CM, Ellis AW (1997) Naming the Snodgrass and Vanderwart pictures: Effects of age of acquisition, frequency, and name agreement. Q J Exp Psychol Sect A 50:560–585.

Bartoli E, Bosking W, Chen Y, Li Y, Sheth SA, Beauchamp MS, Yoshor D, Foster BL (2019) Functionally Distinct Gamma Range Activity Revealed by Stimulus Tuning in Human Visual Cortex. Curr Biol 29:3345–3358.e7.

Benítez-Burraco A, Murphy E (2019) Why brain oscillations are improving our understanding of language. Front Behav Neurosci 13 Available at: https://www.frontiersin.org/articles/10.3389/fnbeh.2019.00190 [Accessed April 9, 2024].

Benjamini Y, Hochberg Y (1995) Controlling the false discovery rate: A practical and powerful approach to multiple testing. J R Stat Soc Ser B Stat Methodol 57:289–300.

Bertrand O, Bohorquez J, Pernier J (1994) Time-frequency digital filtering based on an invertible wavelet transform: an application to evoked potentials. IEEE Trans Biomed Eng 41:77–88.

Binney RJ, Embleton KV, Jefferies E, Parker GJM, Lambon Ralph MA (2010) The ventral and inferolateral aspects of the anterior temporal lobe are crucial in semantic memory: Evidence from a novel direct comparison of distortion-corrected fMRI, rTMS, and semantic dementia. Cereb Cortex 20:2728–2738.

Bozeat S, Lambon Ralph MA, Patterson K, Garrard P, Hodges JR (2000) Non-verbal semantic impairment in semantic dementia. Neuropsychologia 38:1207–1215.

Canolty RT, Edwards E, Dalal SS, Soltani M, Nagarajan SS, Kirsch HE, Berger MS, Barbaro NM, Knight RT (2006) High gamma power is phase-locked to theta oscillations in human neocortex. Science 313:1626–1628.

Canolty RT, Knight RT (2010) The functional role of cross-frequency coupling. Trends Cogn Sci 14:506–515.

Carlyon RP, Deeks JM, McKay CM (2010) The upper limit of temporal pitch for cochlear-implant listeners: Stimulus duration, conditioner pulses, and the number of electrodes stimulated. J Acoust Soc Am 127:1469–1478.

Cervenka MC, Boatman-Reich DF, Ward J, Franaszczuk PJ, Crone NE (2011) Language Mapping in Multilingual Patients: Electrocorticography and Cortical Stimulation During Naming. Front Hum Neurosci 5 Available at: http://journal.frontiersin.org/article/10.3389/fnhum.2011.00013/abstract [Accessed November 4, 2021].

Chan AM, Baker JM, Eskandar E, Schomer D, Ulbert I, Marinkovic K, Cash SS, Halgren E (2011) First-pass selectivity for semantic categories in human anteroventral temporal lobe. J Neurosci 31:18119–18129.

Chaumon M, Bishop DVM, Busch NA (2015) A practical guide to the selection of independent components of the electroencephalogram for artifact correction. J Neurosci Methods 250:47–63.

Chen L, Lambon Ralph MA, Rogers TT (2017) A unified model of human semantic knowledge and its disorders. Nat Hum Behav 1:0039.

Chen Y, Shimotake A, Matsumoto R, Kunieda T, Kikuchi T, Miyamoto S, Fukuyama H, Takahashi R, Ikeda A, Lambon Ralph MA (2016) The ‘when’ and ‘where’ of semantic coding in the anterior temporal lobe: Temporal representational similarity analysis of electrocorticogram data. Cortex 79:1–13.

Clarke A (2020) Dynamic activity patterns in the anterior temporal lobe represents object semantics. Cogn Neurosci 11:111–121.

Clarke A, Devereux BJ, Tyler LK (2018) Oscillatory dynamics of perceptual to conceptual transformations in the ventral visual pathway. J Cogn Neurosci 30:1590–1605.

Cohen MX (2014) Analyzing Neural Time Series Data: Theory and Practice. MIT Press.

Cox CR (2016) Testing neurocognitive predictions of the hub-and-spoke model of semantic memory with network representational similarity analysis.

Cox CR, Rogers TT (2021) Finding distributed needles in neural haystacks. J Neurosci 41:1019–1032.

Cox CR, Rogers TT, Shimotake A, Kikuchi T, Kunieda T, Miyamoto S, Takahashi R, Matsumoto R, Ikeda A, Lambon Ralph MA (2024) Representational similarity learning reveals a graded multidimensional semantic space in the human anterior temporal cortex. Imaging Neurosci 2:1–22.

Crone NE, Hao L (2002) Functional dynamics of spoken and signed word production: A case study using electrocorticographic spectral analysis. Aphasiology 16:903–927.

Crone NE, Hao L, Hart J, Boatman D, Lesser RP, Irizarry R, Gordon B (2001) Electrocorticographic gamma activity during word production in spoken and sign language. Neurology 57:2045–2053.

Delorme A (2023) EEG is better left alone. Sci Rep 13:2372.

Delorme A, Makeig S (2004) EEGLAB: an open source toolbox for analysis of single-trial EEG dynamics including independent component analysis. J Neurosci Methods 134:9–21.

Devlin JT, Russell RP, Davis MH, Price CJ, Wilson J, Moss HE, Matthews PM, Tyler LK (2000) Susceptibility-induced loss of signal: Comparing PET and fMRI on a semantic task. NeuroImage 11:589–600.

Dilkina K, Lambon Ralph MA (2012) Conceptual structure within and between modalities. Front Hum Neurosci 6 Available at: http://journal.frontiersin.org/article/10.3389/fnhum.2012.00333/abstract [Accessed January 10, 2022].

Edwards E, Nagarajan SS, Dalal SS, Canolty RT, Kirsch HE, Barbaro NM, Knight RT (2010) Spatiotemporal imaging of cortical activation during verb generation and picture naming. NeuroImage 50:291–301.

Florez CM, McGinn RJ, Lukankin V, Marwa I, Sugumar S, Dian J, Hazrati L-N, Carlen PL, Zhang L, Valiante TA (2015) In Vitro Recordings of Human Neocortical Oscillations. Cereb Cortex 25:578–597.

Forseth KJ, Kadipasaoglu CM, Conner CR, Hickok G, Knight RT, Tandon N (2018) A lexical semantic hub for heteromodal naming in middle fusiform gyrus. Brain 141:2112–2126.

Frisby SL, Cox CR, Halai AD, Lambon Ralph MA, Rogers TT (2026a) Comparative multivariate decoding adjudicates theories of semantic representation in the anterior temporal lobes and the rest of the cortex. :2025.08.22.671718 Available at: https://www.biorxiv.org/content/10.1101/2025.08.22.671718v2 [Accessed April 27, 2026].

Frisby SL, Halai AD, Cox CR, Clarke A, Shimotake A, Kikuchi T, Kuneida T, Arakawa Y, Takahashi R, Ikeda A, Matsumoto R, Rogers TT, Lambon Ralph MA (2026b) All spectral frequencies of neural activity reveal semantic representation in the human anterior ventral temporal cortex. Imaging Neurosci 4:IMAG.a.1201.

Frisby SL, Halai AD, Cox CR, Lambon Ralph MA, Rogers TT (2023) Decoding semantic representations in mind and brain. Trends Cogn Sci 27:258–281.

Giallanza T, Campbell D, Cohen JD, Rogers TT (2025) An integrated model of semantics and control. Psychol Rev 132:1128–1177.

Halai AD, Welbourne SR, Embleton K, Parkes LM (2014) A comparison of dual gradient-echo and spin-echo fMRI of the inferior temporal lobe. Hum Brain Mapp 35:4118–4128.

Halgren E, Kaestner E, Marinkovic K, Cash SS, Wang C, Schomer DL, Madsen JR, Ulbert I (2015) Laminar profile of spontaneous and evoked theta: Rhythmic modulation of cortical processing during word integration. Neuropsychologia 76:108–124.

Hermes D, Miller KJ, Vansteensel MJ, Edwards E, Ferrier CH, Bleichner MG, van Rijen PC, Aarnoutse EJ, Ramsey NF (2014) Cortical theta wanes for language. NeuroImage 85:738–748.

Heusser AC, Poeppel D, Ezzyat Y, Davachi L (2016) Episodic sequence memory is supported by a theta–gamma phase code. Nat Neurosci 19:1374–1380.

Hodges JR, Patterson K (2007) Semantic dementia: a unique clinicopathological syndrome. Lancet Neurol 6:1004–1014.

Hodges JR, Patterson KE, Oxbury S, Funnell E (1992) Semantic dementia: Progressive fluent aphasia with temporal lobe atrophy. Brain 115:1783–1806.

Jackson RL (2021) The neural correlates of semantic control revisited. NeuroImage 224:117444.

Jackson RL, Rogers TT, Lambon Ralph MA (2021) Reverse-engineering the cortical architecture for controlled semantic cognition. Nat Hum Behav 5:774–786.

Jenkinson M, Beckmann CF, Behrens TEJ, Woolrich MW, Smith SM (2012) FSL. NeuroImage 62:782–790.

Klimesch W (2012) Alpha-band oscillations, attention, and controlled access to stored information. Trends Cogn Sci 16:606–617.

Kojima K, Brown EC, Matsuzaki N, Rothermel R, Fuerst D, Shah A, Mittal S, Sood S, Asano E (2013) Gamma activity modulated by picture and auditory naming tasks: Intracranial recording in patients with focal epilepsy. Clin Neurophysiol 124:1737–1744.

Kopell N, Kramer MA, Malerba P, Whittington MA (2010) Are Different Rhythms Good for Different Functions? Front Hum Neurosci 4 Available at:

https://www.frontiersin.org/articles/10.3389/fnhum.2010.00187 [Accessed April 9, 2024].

Kriegeskorte N, Mur M, Bandettini P (2008a) Representational similarity analysis – connecting the branches of systems neuroscience. Front Syst Neurosci 2 Available at: http://journal.frontiersin.org/article/10.3389/neuro.06.004.2008/abstract [Accessed May 17, 2021].

Kriegeskorte N, Mur M, Ruff DA, Kiani R, Bodurka J, Esteky H, Tanaka K, Bandettini PA (2008b) Matching categorical object representations in inferior temporal cortex of man and monkey. Neuron 60:1126–1141.

Lachaux J-P, Axmacher N, Mormann F, Halgren E, Crone NE (2012) High-frequency neural activity and human cognition: Past, present and possible future of intracranial EEG research. Prog Neurobiol 98:279–301.

Lambon Ralph MA, Jefferies E, Patterson K, Rogers TT (2017) The neural and computational bases of semantic cognition. Nat Rev Neurosci 18:42–55.

Lambon Ralph MA, Patterson K (2008) Generalization and differentiation in semantic memory. Ann N Y Acad Sci 1124:61–76.

Lambon Ralph MA, Sage K, Jones RW, Mayberry EJ (2010) Coherent concepts are computed in the anterior temporal lobes. Proc Natl Acad Sci 107:2717–2722.

Lisman J (2005) The theta/gamma discrete phase code occuring during the hippocampal phase precession may be a more general brain coding scheme. Hippocampus 15:913–922.

Lisman J, Jensen O (2013) The theta-gamma neural code. Neuron 77:1002–1016.

Lüders H, Lesser RP, Hahn J, Dinner DS, Morris HH, Wylie E, Godoy J (1991) Basal temporal language area. Brain 114:743–754.

Marko M, Cimrová B, Riečanský I (2019) Neural theta oscillations support semantic memory retrieval. Sci Rep 9:17667.

Matoba K, Matsumoto R, Shimotake A, Nakae T, Imamura H, Togo M, Yamao Y, Usami K, Kikuchi T, Yoshida K, Matsuhashi M, Kunieda T, Miyamoto S, Takahashi R, Ikeda A (2024) Basal temporal language area revisited in Japanese language with a language function density map. Cereb Cortex 34:bhae218.

Mervis CB, Rosch E (1981) Categorization of natural objects. Annu Rev Psychol 32:89–115.

Miller KJ (2010) Broadband spectral change: Evidence for a macroscale correlate of population firing rate? J Neurosci 30:6477–6479.

Mitra P, Bokil H (2007) Observed Brain Dynamics. Oxford University Press.

Mollo G, Cornelissen PL, Millman RE, Ellis AW, Jefferies E (2017) Oscillatory dynamics supporting semantic cognition: MEG evidence for the contribution of the anterior temporal lobe hub and modality-specific spokes Urgesi C, ed. PLOS ONE 12:e0169269.

Morrison CM, Chappell TD, Ellis AW (1997) Age of acquisition norms for a large set of object names and their relation to adult estimates and other variables. Q J Exp Psychol Sect A 50:528–559.

Murphy E (2024) ROSE: A neurocomputational architecture for syntax. J Neurolinguistics 70:101180.

Murphy E, Woolnough O, Morse CW, Scherschligt X, Tandon N (2026) Frontotemporal network interactions causally support rapid concreteness judgments during reading. PLOS Biol 24:e3003723.

Nakai Y, Jeong J, Brown EC, Rothermel R, Kojima K, Kambara T, Shah A, Mittal S, Sood S, Asano E (2017) Three- and four-dimensional mapping of speech and language in patients with epilepsy. Brain 140:1351–1370.

Nakai Y, Sugiura A, Brown EC, Sonoda M, Jeong J, Rothermel R, Luat AF, Sood S, Asano E (2019) Four-dimensional functional cortical maps of visual and auditory language: Intracranial recording. Epilepsia 60:255–267.

Oswal U, Cox CR, Lambon Ralph MA, Rogers TT, Nowak RD (2016) Representational Similarity Learning with Application to Brain Networks. In: Proceedings of The 33rd International Conference on Machine Learning, pp 1041–1049. PMLR. Available at: https://proceedings.mlr.press/v48/oswal16.html [Accessed August 26, 2024].

Patterson K, Nestor PJ, Rogers TT (2007) Where do you know what you know? The representation of semantic knowledge in the human brain. Nat Rev Neurosci 8:976–987.

Pobric G, Jefferies E, Lambon Ralph MA (2007) Anterior temporal lobes mediate semantic representation: Mimicking semantic dementia by using rTMS in normal participants. Proc Natl Acad Sci 104:20137–20141.

Pobric G, Jefferies E, Lambon Ralph MA (2010a) Category-specific versus category-general semantic impairment induced by transcranial magnetic stimulation. Curr Biol 20:964–968.

Pobric G, Jefferies E, Lambon Ralph MA (2010b) Amodal semantic representations depend on both anterior temporal lobes: Evidence from repetitive transcranial magnetic stimulation. Neuropsychologia 48:1336–1342.

Proix T, Delgado Saa J, Christen A, Martin S, Pasley BN, Knight RT, Tian X, Poeppel D, Doyle WK, Devinsky O, Arnal LH, Mégevand P, Giraud A-L (2022) Imagined speech can be decoded from low- and cross-frequency intracranial EEG features. Nat Commun 13:48.

Quiroga RQ, Kreiman G, Koch C, Fried I (2008) Sparse but not ‘Grandmother-cell’ coding in the medial temporal lobe. Trends Cogn Sci 12:87–91.

Rogers TT, Cox CR, Lu Q, Shimotake A, Kikuchi T, Kunieda T, Miyamoto S, Takahashi R, Ikeda A, Matsumoto R, Lambon Ralph MA (2021) Evidence for a deep, distributed and dynamic code for animacy in human ventral anterior temporal cortex. eLife 10:e66276.

Rogers TT, Lambon Ralph MA, Garrard P, Bozeat S, McClelland JL, Hodges JR, Patterson K (2004) Structure and deterioration of semantic memory: a neuropsychological and computational investigation. Psychol Rev 111:205–235.

Rogers TT, McClelland JL (2004) Semantic Cognition: A Parallel Distributed Processing Approach. MIT Press.

Rogers TT, Patterson K (2007) Object categorization: Reversals and explanations of the basic-level advantage. J Exp Psychol Gen 136:451–469.

Rupp K, Roos M, Milsap G, Caceres C, Ratto C, Chevillet M, Crone NE, Wolmetz M (2017) Semantic attributes are encoded in human electrocorticographic signals during visual object recognition. NeuroImage 148:318–329.

Sato N, Matsumoto R, Shimotake A, Matsuhashi M, Otani M, Kikuchi T, Kunieda T, Mizuhara H, Miyamoto S, Takahashi R, Ikeda A (2021) Frequency-dependent cortical interactions during semantic processing: an electrocorticogram cross-spectrum analysis using a semantic space model. Cereb Cortex 31:4329–4339.

Sauseng P, Klimesch W (2008) What does phase information of oscillatory brain activity tell us about cognitive processes? Neurosci Biobehav Rev 32:1001–1013.

Sederberg PB, Kahana MJ, Howard MW, Donner EJ, Madsen JR (2003) Theta and gamma oscillations during encoding predict subsequent recall. J Neurosci 23:10809–10814.

Sevgili I (2023) Unravelling Spiral Ganglion Neuron Electrophysiology: Heterogeneity, Gene Therapy, and In-Vitro Testing Models. Available at: https://www.repository.cam.ac.uk/handle/1810/362231 [Accessed May 1, 2026].

Shimotake A, Matsumoto R, Ueno T, Kunieda T, Saito S, Hoffman P, Kikuchi T, Fukuyama H, Miyamoto S, Takahashi R, Ikeda A, Lambon Ralph MA (2015) Direct exploration of the role of the ventral anterior temporal lobe in semantic memory: cortical stimulation and local field potential evidence from subdural grid electrodes. Cereb Cortex 25:3802–3817.

Smith SM, Jenkinson M, Woolrich MW, Beckmann CF, Behrens TEJ, Johansen-Berg H, Bannister PR, De Luca M, Drobnjak I, Flitney DE, Niazy RK, Saunders J, Vickers J, Zhang Y, De Stefano N, Brady JM, Matthews PM (2004) Advances in functional and structural MR image analysis and implementation as FSL. NeuroImage 23:S208–S219.

Snyder KM, Forseth KJ, Donos C, Rollo PS, Fischer-Baum S, Breier J, Tandon N (2023) Critical role of the ventral temporal lobe in naming. Epilepsia 64:1200–1213.

Solomon EA, Lega BC, Sperling MR, Kahana MJ (2019) Hippocampal theta codes for distances in semantic and temporal spaces. Proc Natl Acad Sci 116:24343–24352.

Spiers HJ (2020) The hippocampal cognitive map: One space or many? Trends Cogn Sci 24:168–170.

Tanji K (2005) High-frequency -band activity in the basal temporal cortex during picture-naming and lexical-decision tasks. J Neurosci 25:3287–3293.

Visser M, Jefferies E, Lambon Ralph MA (2010) Semantic processing in the anterior temporal lobes: a meta-analysis of the functional neuroimaging literature. J Cogn Neurosci 22:1083–1094.

Wang W, Degenhart AD, Sudre GP, Pomerleau DA, Tyler-Kabara EC (2011) Decoding semantic information from human electrocorticographic (ECoG) signals. Annu Int Conf IEEE Eng Med Biol Soc IEEE Eng Med Biol Soc Annu Int Conf 2011:6294–6298.

Watrous AJ, Deuker L, Fell J, Axmacher N (2015) Phase-amplitude coupling supports phase coding in human ECoG. eLife 4:e07886.

